# Object dimensions underlying food representations in visual cortex

**DOI:** 10.64898/2026.08.24.746622

**Authors:** Davide Cortinovis, Giulia Orlandi, Lotte van Campenhout, Stefania Bracci

**Affiliations:** Center for Mind/Brain Sciences (CIMeC), University of Trento, Rovereto, 38068, Italy; Brain and Cognition, Faculty of Psychology and Educational Sciences, KU Leuven, 3000, Leuven, Belgium

## Abstract

Recent work has revealed two food-selective areas in the lateral and ventromedial occipitotemporal cortex (OTC). These studies have shown that food selectivity in these regions cannot be explained by mid-level features like shape, texture, or colour but differences in their representational content remain unclear. Across two fMRI experiments conducted in the same group of participants, we characterized the dimensions underlying food representations in lateral and ventral OTC by examining the contribution of action-related object properties, such as manipulability, relevant to object interaction, and visual features, such as colour and ensemble statistics, relevant to object recognition. Our results reveal a clear dissociation between lateral and ventral OTC, indicating that food representations in these regions reflect distinct computational constraints. In lateral OTC, food representations were primarily associated with action-related properties shared between food and other graspable objects, whereas in ventral OTC, food representations were sensitive to surface object properties, such as colour and ensemble configuration. Consistent with this distinction, lateral OTC showed greater sensitivity to individual objects against distinctive background and responded equally to colour and greyscale stimuli, while ventral OTC exhibited greater sensitivity to coloured stimuli and ensembles with no distinctive background. Finally, topographic artificial neural networks implementing architectural constraints meant to capture OTC spatial organization similarly exhibited two dissociable clusters of food-selective units based on sensitivity to ensemble statistics. Together, these findings suggest that lateral food representations reflect action-relevant properties shared with other manipulable objects, whereas ventral food representations arise from surface-based visual features critical for food identification.

**Significant Statement:** Two areas of human occipitotemporal cortex (OTC) respond selectively to food, yet whether they serve distinct computational roles remains unclear. Across two fMRI experiments using controlled stimulus sets, we show that these areas support different visual and non-visual goals. Lateral OTC represents food together with other manipulable objects, reflecting action-related properties relevant for interacting with objects, whereas ventral OTC is driven by surface properties, particularly colour and ensemble configuration, relevant for food identification. Topographic neural networks reproduced two food clusters dissociable by ensemble statistics, but not the colour-or action-related organization observed in cortex. These findings indicate that food-selective areas reflect the behaviourally relevant feature dimensions computed by the pathway in which they reside, rather than category membership per se.

## Introduction

The occipitotemporal cortex (OTC) is organized into areas that respond selectively to ecologically relevant object categories, including faces, hands, tools, and scenes (Kanwisher, 2010; Grill-Spector and Weiner, 2014; Arcaro and Livingstone, 2024). These areas are embedded within larger-scale maps representing dimensions such as eccentricity (Hasson et al., 2003), curvature (Yue et al., 2020), animacy (Kriegeskorte et al., 2008a), real-world size (Konkle and Oliva, 2012), and action (Cortinovis et al., 2025a). For example, hand-and tool-selective areas are adjacent and partially overlapping, reflecting shared sensitivity to action-related properties (Bracci et al., 2012). A similar nested organization emerges in Topographic Deep Artificial Neural Networks (TDANNs), which impose correlations among neighbouring units to simulate OTC spatial organization (Margalit et al., 2024; Deb et al., 2025; Lu et al., 2025).

Recent research has focused on another object category: food. Food is among the most behaviourally-salient categories, engaging visual cortex alongside taste-and reward-related regions such as the insula and orbitofrontal cortex (Simmons et al., 2005; Avery et al., 2021; van der Laan et al., 2011). Studies identified robust food-selective responses in ventral visual cortex (Khosla et al., 2022; Jain et al., 2023; Pennock et al., 2023) comparable to classic category-selective responses, reporting two food-selective “stripes”: one in ventral and one in lateral OTC, which responses could not be explained by low-or mid-level features such as curvature or texture, providing evidence for food-category selectivity.

Despite this progress, the extent to which these two food-selective clusters reflect distinct representational or computational profiles remains unclear. One study reported a partial dissociation based on image content using PCA (close-up food versus food-related scenes; Jain et al., 2023), but most analyses detected no differences in feature sensitivity. In light of recent accounts proposing a division of labour between ventral and lateral OTC in object recognition and action processing, respectively (Lingnau and Downing, 2015; Wurm and Caramazza, 2022), food is ideal for testing whether these regions contribute differently matching their computational goals.

Several properties may drive food selectivity (reviewed in Henderson et al., 2025). Here we focus on three: two visual (colour and ensemble statistics) and one action-related (graspability/manipulability). Colour is an obvious candidate: it conveys food-relevant information such as ripeness and calorie content (Foroni et al., 2016) and supports rapid recognition (Sato, 2021). Colour responses have been documented in primate visual cortex (Conway et al., 2007; Lafer-Sousa et al., 2016), including regions near food-selective areas, with colour-biased patches argued to be food-selective (Pennock et al., 2023). A less explored property is ensemble configuration: food often appears as collections of similar elements (e.g., peas, grapes). While ensemble perception is well studied behaviourally (Alvarez, 2011; Whitney and Yamanashi Leib, 2018), its neural basis remains unclear; some evidence points to regions near the parahippocampal place area (Cant and Xu, 2012), potentially overlapping with ventral food-selective cortex. Finally, food is inherently action-related, as it must be manipulated before consumption, and a recent study reported overlapping food and tool activations in ventral and lateral OTC, suggesting graspability/manipulability as a driver of food representations (Ritchie et al., 2024).

We therefore hypothesized that food selectivity reflects two dimensions: (1) a visual dimension capturing surface-level properties such as colour and ensemble statistics, relevant for object recognition and driving responses in ventral OTC; and (2) an action-related dimension defined by object affordances, relevant for action processing and driving responses in lateral OTC. Across two fMRI experiments, we observed a robust dissociation between ventral and lateral food areas. Ventral clusters were primarily tuned to surface properties, especially colour and, to a lesser extent, ensemble configuration, whereas the left lateral cluster was more strongly associated with manipulability properties shared with other objects, consistent with their different computational role. Applying the same analyses to TDANNs revealed two food-responsive clusters likewise dissociable by sensitivity to ensemble statistics, partially resembling cortical organization. However, no comparable organization based on colour or action-related properties emerged in the model.

## Methods

### fMRI experimental design

#### Participants

A total of 20 participants took part in both fMRI experiments: the *category experiment* and the *surface properties experiment*. Two participants were excluded due to excessive head motion (exceeding one voxel in translation or rotation), resulting in a final sample of 18 participants (9 females, sex self-reported; mean age = 24 years, SD = 3.3). All participants were right-handed, had normal or corrected-to-normal vision, and no history of neurological disorders. Written informed consent was obtained from all participants, and the experimental procedures were approved by the Ethics Committee of the University of Trento.

#### Stimuli

The *category experiment* included 8 categories. The set comprises faces, headless bodies, hands, food, tools, manipulable objects, scenes, and meaningless shapes. Food category included both graspable food (e.g., hot-dog, sandwich) and non-directly graspable food (e.g., lasagna). Tools were defined as hand-held objects that are typically used to physically and directly act on another object or surface (e.g., pliers, knife), whereas manipulable objects were defined as objects that can be grasped but are not usually used as action-effectors (e.g., bedside lamp, book; see Bracci and Peelen, 2013). The two object categories were matched for overall shape and orientation. Each category included 48 greyscale (400×400) images with a white background. Part of the images were used in Matić et al. (2020); part of the images of food were selected among the localizers developed by Jain et al., (2023) and Ritchie et al. (2024); meaningless shapes were taken from Op de Beeck et al. (2006); all other images were generated from images downloaded from the internet.

The *surface properties experiment* included 8 conditions, varying along three orthogonal dimensions: 1) category (food vs objects), 2) colour (coloured vs. greyscale), and 3) ensemble/configuration (images were either presented as food/object ensembles, such as a group of apples, or as the corresponding single object, such as a single apple on a naturalistic background). The stimuli were selected to minimize potential confounds between categories: for instance, we ensured that the inanimate object categories contained stimuli that were as colourful as food images, and we selected food stimuli to contain various types of colours (both warm and cooler colours). All images were sourced through internet searches. Examples of stimuli for both sets can be visualized in Figure 1.

**Figure 1.**
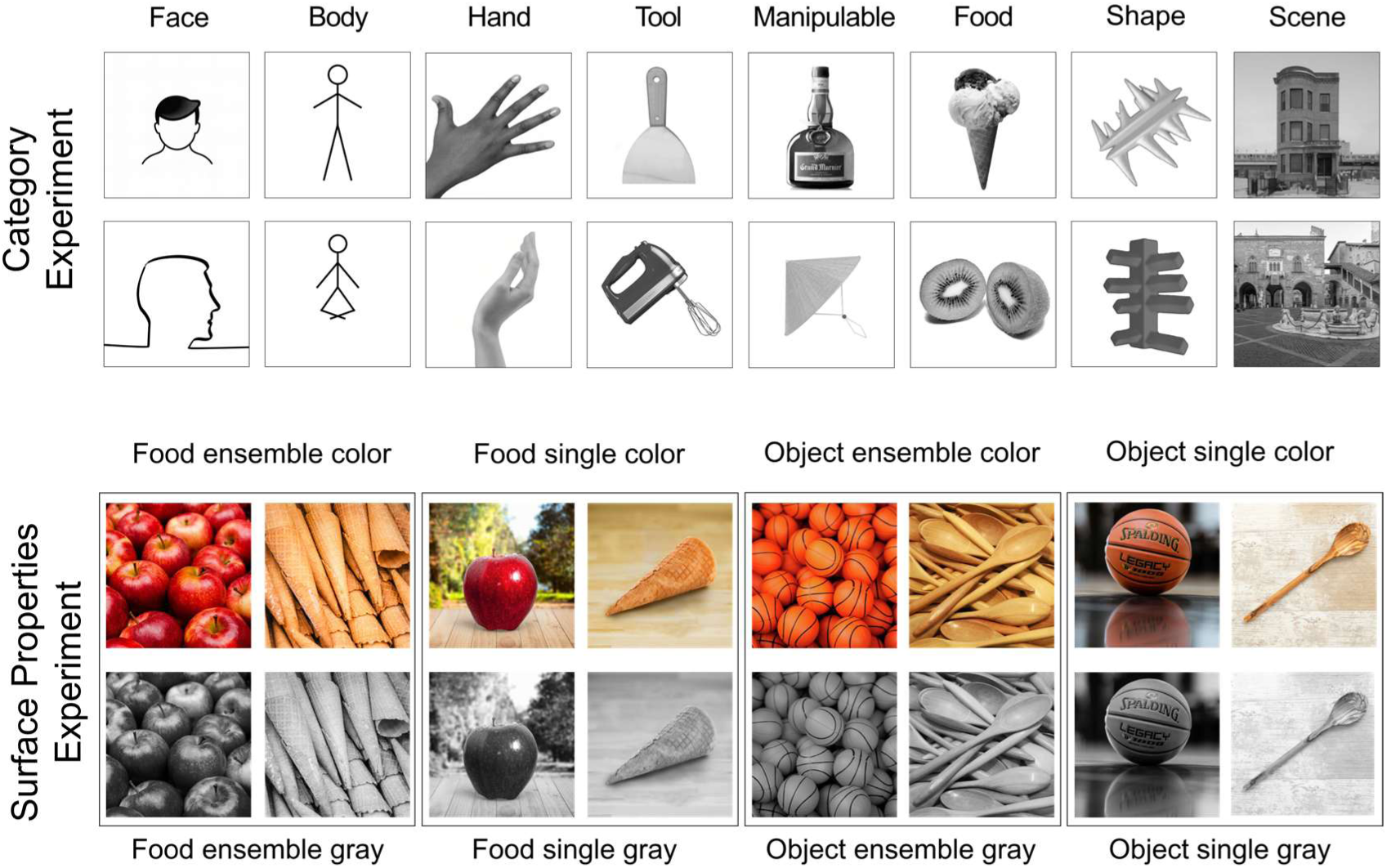
Stimulus sets. Examples of images from the *category experiment* (above) and the *surface properties experiment* (below).

#### Scanning procedure

The fMRI study was conducted over two sessions on separate days within a week. In the first session we collected the data for the *category experiment* (6 runs per participant) and in the second session, we collected data for the *surface properties experiment* (6 runs per participant). The anatomical scan was collected during the first session. The design was the same for both experiments. Each run lasted 336 seconds (168 volumes). Images were presented for 400 ms each, with an inter-stimulus interval (ISI) of 266 ms, organized into 8-second blocks (12 images per block). For each participant and run, a fully randomized sequence of all conditions was repeated 4 times, separated with 16-second fixation blocks, including at the beginning and end of the run. The entire experiment therefore consisted of 24 repetitions of each category block. Stimuli were displayed using the Psychophysics Toolbox (Brainard 1997) with MATLAB (version 2022b; The MathWorks) and projected onto a screen subtending 8 × 8 degrees of visual angle, viewed via a mirror on the head coil. Participants were instructed to maintain fixation on a central cross and press a button whenever the same image appeared twice consecutively within a block (one repeat per block). Behavioural performance was quantified by calculating response accuracy (mean across experiments = 93.7%, SD = 3.1%) and reaction time (RT; mean across experiments = 588.6 ms, SD = 25.8 ms). Accuracy was defined as the proportion of correctly identified target stimuli. Given the rapid presentation rate (400 ms), responses were considered correct if made within two trials after target onset.

#### Imaging parameters

The fMRI data was collected using a 3T Siemens scanner with a 64-channel head coil in the Center for Mind/Brain Sciences at the University of Trento. MRI volumes were collected using echo planar (EPI) T2*-weighted sequence, with repetition time (TR) of 2 s, echo time (TE) of 28 ms, flip angle (FA) of 75°, and field of view of 220 mm. Each volume contained 69 axial slices, covering the whole brain, with matrix size 200×200 mm and 2×2×2 mm voxel size. Slices were acquired with a multiband (multi-slice) sequence, with slice acceleration factor = 3. Anatomical images were acquired using the T1-weighted acquisition and MP-RAGE sequence, with a resolution of 1×1×1 mm.

#### Preprocessing

Preprocessing was performed using the Statistical Parametric Mapping software package (SPM12, Wellcome Trust Centre for Neuroimaging, London) and MATLAB (R2021b, The MathWorks). Functional images underwent the following preprocessing steps: spatial realignment to the first image to correct for head motion, slice-timing correction, coregistration with anatomical images, and spatial smoothing using a Gaussian kernel (4 mm FWHM). When performing group-level analysis, the data was normalized to the Montreal Neurological Institute’s (MNI) ICBM152 template. Prior to preprocessing, we defined participants’ exclusion criteria as follows: runs where head movement exceeded one voxel (in translation or rotation, 2 mm) were excluded; if more than half of the runs (6 total per experiment) were excluded for a given participant, that participant was fully excluded from further analysis. Based on this criterion, two participants were excluded entirely, along with 3 runs from 3 participants (one run each) for the *category experiment* and 4 runs from 3 participants (one run in two participants and two runs in one participant) for the *surface properties experiment*. The preprocessed signal was modelled for each voxel, for each participant, and for each condition using a general linear model (GLM). For both experiments, the GLM included 8 regressors of interest (one per experimental condition) and 6 nuisance regressors (motion correction parameters: x, y, z for translations and rotations). Predictors’ time courses were modelled by convolving the haemodynamic response function (HRF) with a boxcar function.

### Data analysis

#### Whole-brain univariate analysis

First, we performed a group random-effects analysis to visualize the topographic organization of food-selective areas and their spatial relationship with other category-selective responses for both experiments. Each condition in the *category experiment* was contrasted against the average of all the others (e.g., food vs all), whereas each condition in the *surface properties experiment* (category, colour, and configuration) was contrasted against the opposite condition (i.e., food vs. objects; colour vs. grayscale; and ensemble vs. single). Results were thresholded at *p* < .001 uncorrected at the voxel level and *p* < .05 FDR corrected at the cluster level and visualized on a Freesurfer average surface using BrainSurfer (Teghipco, 2023).

#### ROI definition

We identified ROIs at both the individual and at the group level for both studies. First, for both experiments, we identified areas selectively responding to food images on the native volume of each individual. These areas were defined with a contrast of category vs. average of all others, with a threshold of *p* < .001 uncorrected or – if no ROI could be selected at this threshold – p < .01 uncorrected. If necessary (e.g., when the activation formed a contiguous cluster across lateral and ventral OTC or with activations in early visual cortex), ROIs were restricted to a cube of 6 mm width centred on the activation peak. The areas defined in one experiment were used to test functional selectivity in the other experiment and viceversa, thus ensuring independence of the data used for statistical analysis from the ROI definition. Food areas were defined bilaterally and separately for ventral and lateral OTC.

To test overlap between categories, we identified additional ROIs using the same criteria as above. For the *category experiment*, we identified hand and tool areas with a contrast of *p* < .001 uncorrected. Activations for manipulable objects could not be reliably defined at this threshold, so we did not include this category in the overlap analysis. For the *surface properties experiment*, we identified areas selective to colour with a contrast of p < .001 uncorrected or – if no ROI could be selected at this threshold – p < .01 uncorrected. All ROIs were defined bilaterally, except tool-selective areas that exhibited a strong left-lateralization.

To test multivariate patterns for the *category experiment* four ROIs were identified at the group level: VOTC (left and right) and LOTC (left and right). These areas were defined using the Glasser atlas (Glasser et al., 2016) based on a combination of functional and anatomical criteria as follows: first, for VOTC, we selected the following areas: FFC, VVC, VMV1-3, PHA1-3, PIT; for LOTC the following areas: LO1-3, MT/MST, PH, FST, TE1p/TE2p. Then, we included all voxels within those anatomical masks that responded to all categories vs. baseline and survived the threshold at p < .001 uncorrected. These ROIs were used to test multivariate patterns in regions of cortex that include but go beyond food responses.

#### Functional selectivity analyses

For each experiment and each ROI, we first examined functional selectivity. In the *category experiment*, we assessed food selectivity relative to a range of visual categories to test whether food-selective areas are truly specific to food stimuli or also respond to other categories. In the *surface properties experiment*, we evaluate whether food-and object-selective areas respond differentially to surface properties such as colour and ensemble statistics. This analysis also enabled us to explore potential differences in responses between ventral and lateral food-selective clusters. Importantly, by defining ROIs using data from the other experiment, we ensured statistical independence between ROI definition and functional selectivity assessment.

#### Overlap analyses

For both experiments, anatomical overlap between categories (or conditions) was measured using ROIs defined for each subject in native space, following a procedure used in previous studies (e.g. Bracci et al., 2012). Specifically, we calculated the number of voxels in common between two ROIs (e.g., hands and food or colour and food) and divided it by the smaller of the two ROIs. This gives us an index ranging from 0 (no overlap between the two categories) to 1 (full overlap, where the smaller of the two ROIs falls completely within the bigger of the two).

#### Vector-of-ROIs

To investigate the broader topographic organization of object categories and feature dimensions, we conducted a vector-of-ROIs analysis (originally developed by Konkle and Caramazza, 2013). This procedure involves the construction of a series of partially overlapping spheres along a vector. The vector was generated by fitting a spline connecting two points: one medial, around the parahippocampal gyrus, and one dorsal and lateral, around the transverse occipital sulcus. Coordinates for these two points were taken from Konkle and Caramazza (2013). Between these two points, a series of anchor points was defined to constraint the vector to pass through previously known areas responding to specific object categories. The anchor points coordinates were based on published studies on category selectivity and specifically passed through areas responding to faces (Julian et al., 2012), small objects (Konkle and Oliva, 2012), hands and tools (Bracci et al., 2012), and bodies (Julian et al., 2012). In addition to these areas, we also added an anchor point that was based on the coordinates of the food-selective area in the ventral OTC, to make sure that the vector passed through the relevant regions showing food preference. As before, food coordinates were defined independently: for the *category experiment* data, we used the food peak coordinates derived from the *surface properties experiment*, and viceversa. Along the vector, a series of spheres of 5 mm, spaced 3 mm between each other, was generated. We extracted the beta values for both experiments and plotted the activation for each category and each condition in each of the sphere and analysed their functional profile.

#### Representational similarity analysis

To investigate the representational space underlying food-selective areas we adopted a Representational Similarity Analysis (RSA) approach (Kriegeskorte et al., 2008b). For each experiment, we computed correlation matrices by calculating the pairwise correlations of all conditions based on the group-derived ROIs, leading to 8×8 matrices for bilateral ventral and lateral OTC.

In the *category experiment*, the resulting similarity matrices were used to test the role of action properties in shaping the organization of food and other categories in visual cortex. To this aim, we computed two indices: the food index and the hand index. The food index measured the similarity between food and other categories that share manipulability properties (tools, manipulable objects, and hands). For each participant, we calculated the pairwise correlations between food and each of the other categories (food–tool, food–manipulable, and food–hand) and subtracted the average of the two remaining correlations; that is food-tool – avg(food-hand, food-manipulable), food-manipulable – avg(food-hand, food-tool), and food-hand – avg(food-tool, food-manipulable). The hand index was intended to measure the action effector properties shared between hands and other object categories (food, tools, manipulable objects). As above, for each participant, we calculated the pairwise correlations between hand and each of the other categories (hand–tool, hand–manipulable, and hand-food) and subtracted the average of the two remaining correlations; that is hand-tool – avg(hand-food, hand-manipulable), hand-manipulable – avg(hand-tool, hand-food), and hand-food – avg(hand-tool, hand-manipulable).

In the *surface properties experiment*, to investigate the role of colour and ensemble statistics in shaping food-related representational content, we correlated the obtained matrices with two theoretical models that captures the colour (coloured vs greyscale images) and ensemble (groups of objects vs single object) dimensions. We then ran semi-partial correlations between the models and each ROI and tested each model’s ability to explain independent portions of variance in the ROI’s representational space. By design, the two models were orthogonal (r = 0). The lower bound of the noise ceiling was calculated by correlating each participant matrix with the group average matrix iteratively excluding the participant being correlated.

#### Statistical analysis

For the functional selectivity analyses, beta values for each category (or condition) were extracted from each participant’s ROIs and compared using two-tailed t-tests. For each region, responses were tested against baseline with one-sample t-tests (Bonferroni corrected at *p* < .006, *n* = 8 comparisons) and across categories with pairwise paired t-tests (Bonferroni corrected at *p* < .007, *n* = 7 comparisons). For the multivariate analyses, Fisher-transformed correlation values were tested with repeated-measures ANOVAs and post-hoc two-tailed paired t-tests, Bonferroni corrected when relevant to account for multiple comparisons.

### Topographic Artificial Neural Networks

We adopted a recent computational model developed to capture the spatial organization of ventral temporal cortex – the TDANN (Margalit et al., 2024) – to test its ability in replicating the spatial and functional organization of food responses in visual cortex. Here, we briefly describe its main characteristics.

The model is based on a ResNet-18 architecture; it includes nine topographic layers (meant to capture the entire hierarchical organization of the ventral visual stream); each topographic layer is a grid of units, and each unit is assigned to a specific coordinate before training based on pre-set optimizations such as coarse retinotopic mapping. During training, the model concurrently optimizes two loss functions: a task loss, where the model is trained with a self-supervised contrastive learning task (SimCLR; Chen et al., 2020) on ImageNet (Deng et al., 2009), and the spatial loss, where units that are neighbouring in the layer must have correlated firing patterns; the strength of correlation is determined by a parameter called alpha, that is set at 0.25. We performed the same analysis as done in visual cortex. We focus on the last “VTC-like” topographic layer, meant to capture the organization of high-level ventral visual cortex, and we test five random initializations of the network’s weights.

More specifically, we tested the topographic organization and selectivity profile of the five different random initializations of the network in response to our eight object categories (in the *category experiment*) and our three dimensions (in *the surface properties experiment*). First, we tested the spatial clustering of units selective for the different object categories\dimensions within the simulated physical cortical space in the VTC-like layer. Then, for more precise functional selectivity analysis, we identify spatially contiguous clusters of food-selective units in the VTC-like layer by adopting a patch-based clustering approach using the DBSCAN algorithm (Ester et al., 1996). Food selectivity was first quantified in each unit using a t-statistic comparing responses to food images against the other categories. Units exceeding a fixed selectivity threshold (t > 5.0) were considered supra-threshold candidates. DBSCAN clustering was then applied to the spatial coordinates of these units to identify contiguous patches, with a fixed neighbourhood radius of 0.3. For each model instance, we analysed the two largest significant food-selective patches. The spatial clustering and functional selectivity analyses were also computed for each stimulus set within these food-selective clusters.

## Results

In this study, we investigated the potential differences in how food is represented in the ventral and lateral occipitotemporal cortex (OTC). Specifically, the aim was to test if these regions exhibit differential sensitivity to action-related versus visual surface properties reflecting the functional distinction between the ventral and lateral pathways. Two fMRI experiments in the same group of participants were conducted. In the *category experiment*, participants viewed stimuli from eight object categories to examine the topographic organization of food and other object representations, specifically those characterized by action-related properties, such as graspability/manipulability (shared with tools and manipulable objects) and end-effector properties (shared by hands and tools but not by food). In the *surface properties experiment*, participants were presented with images of food and objects that varied orthogonally along two dimensions: colour (coloured vs. grayscale images), and configuration (ensemble vs. single objects), to test how surface properties modulate responses in food areas. Example stimuli from both experiments are shown in Figure 1.

### The topographic organization of food areas in ventral and lateral OTC

We first mapped food activations and examined their spatial organization relative to other category-selective areas at the group level (p < .001 uncorrected at the voxel level and p <.05 FDR-corrected at the cluster level) across both ventral and lateral OTC (Figure 2). Our results replicated and extended recent reports (Khosla et al., 2022; Jain et al., 2023; Pennock et al., 2023) highlighting two separate food clusters: one in ventral OTC, located between face-and scene-selective areas, and one in lateral OTC.

**Figure 2.**
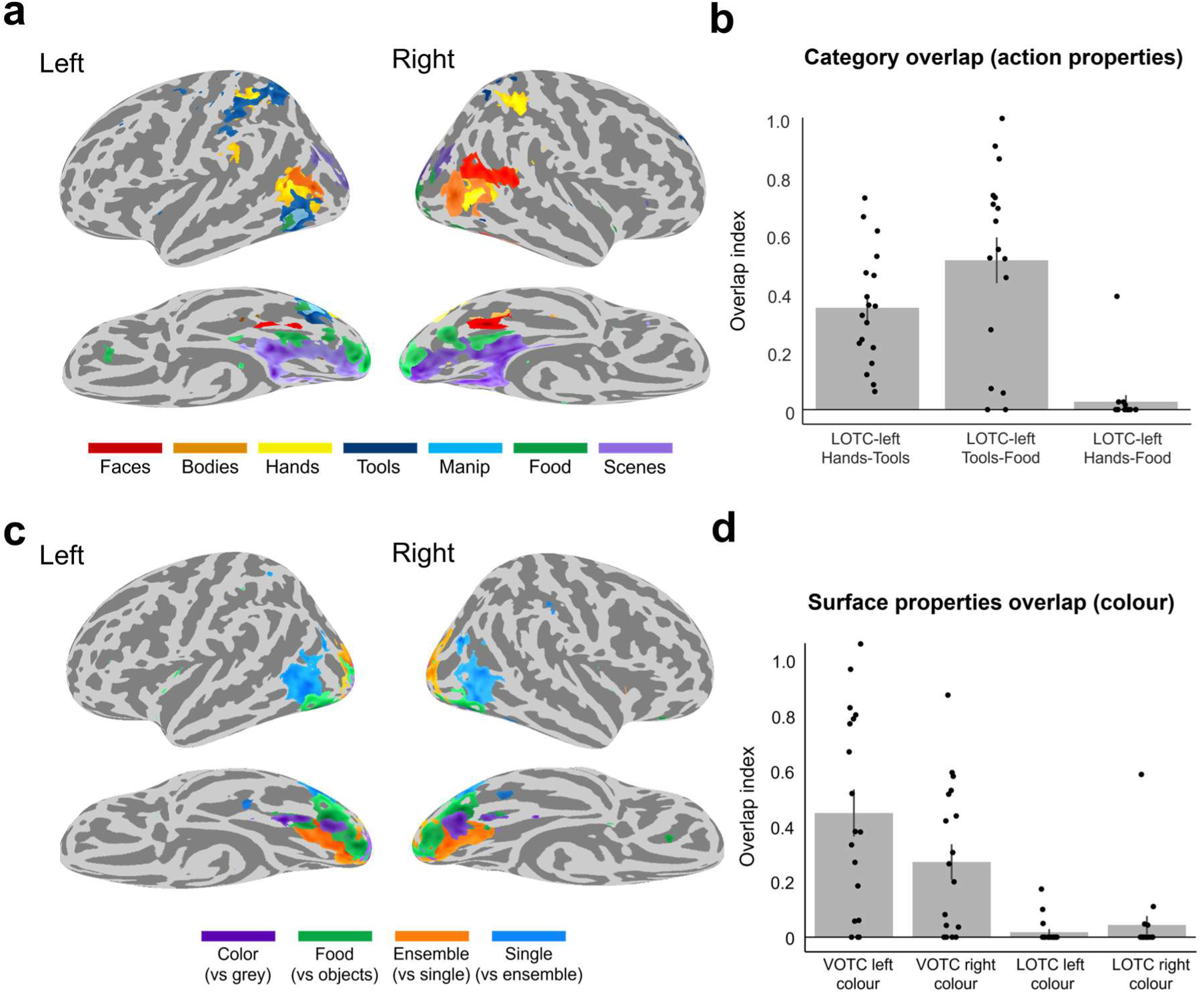
The topography of food areas in occipitotemporal cortex. a) Group-level whole-brain results for the *category experiment*. Response for each category (vs. all) was visualized on a freesurfer average brain surface using BrainSurfer (Teghipco, 2023), with a threshold of t > 3.5. b) Category overlap (action). Overlap index is plotted for each pairwise overlap among three categories (hands, food, tools) in left LOTC. c) Group-level whole-brain results for the *surface properties experiment*. Same visualization as a). d) Surface properties overlap (colour). Overlap index is plotted for each pairwise overlap among two conditions (colour and food) and four areas (VOTC and LOTC, left and right hemisphere). Error bars represent ± SEM across subjects (n = 18 participants). Each data point represents the overlap value from one subject.

In lateral OTC food images activate regions, in the left hemisphere, that respond to tools and other manipulable objects, anterior to – but nonoverlapping with – hand representations. Tool activations extended more posteriorly, overlapping partially with the hand cluster, while the body activation (smaller in the left than in the right hemisphere) was even more posterior, overlapping with hand-but not tool-related clusters. This result is consistent with our recent proposal (Cortinovis et al., 2025a) that in lateral OTC object representations are organized according to an action-related principle progressing from effector-specific to graspable/manipulable properties along a posterior-to-anterior axis, therefore predicting that food clusters should emerge in locations associated with graspable (e.g., tools and manipulable objects), but not action effector representations (e.g., hands, but see Ritchie et al., 2024). In line with this prediction, an overlap analysis (Figure 2b) performed on single-subject native space maps (see methods) confirmed that in lateral OTC, food voxels overlap substantially with tool voxels (score = 0.51, p < .001) but not with hand voxels (score = 0.03, p = .13). In contrast, hand and tool areas showed substantial overlap (score = 0.35, p < .001).

Next, to visualize the relation between food representations and object surface properties, using data from the *surface properties experiment*, we mapped each property – colour and ensemble statistics – and assessed their relative spatial arrangement with food areas within ventral and lateral OTC (Figure 2c). The food > object contrast confirmed the results observed in the *category experiment*, revealing two food clusters: one in ventral and one in lateral OTC. As for surface properties, the configuration contrast (ensembles > single objects) produced robust activation in bilateral early visual cortex extending more anteriorly to ventral OTC, whereas the reverse contrast (single objects > ensembles) yielded activations in lateral OTC. While the contrast (grayscale > colour) did not reveal any activation, the colour contrast (colour > grayscale), revealed two main colour-selective clusters in ventral OTC, in line with prior findings (Lafer-Sousa et al., 2016; Pennock et al., 2023): a posterior cluster likely corresponding to classically defined area V4, and a more anterior cluster in close proximity to the ventral food cluster. The overlap analysis (Figure 2d) performed on single-subject native space maps (see methods) quantified these spatial relationships. The ventral food cluster exhibited a substantial overlap with the colour cluster in both hemispheres (left: overlap score = 0.42, p < .001; right: overlap score = 0.26, p < .001), whereas the lateral food cluster did not (left: overlap score = 0.02, p = 0.06; right: overlap score = 0.04, p = 0.1), consistent with the known ventral–lateral dissociation of colour responses (Brouwer and Heeger, 2009).

Together, these results suggest a dissociation between ventral and lateral OTC food organization: lateral food clusters might reflect graspable properties shared by food, tools, and other manipulable objects, whereas ventral food clusters represent surface properties.

### Functional selectivity analyses reveal a dissociation between ventral and lateral OTC food representations

To characterize the functional tuning of ventral and lateral OTC food areas, ROIs were identified independently in each participant’s native volume (see methods) and used to measure responses to the stimulus categories within each study. Across both studies, results confirm a functional dissociation between ventral and lateral OTC food areas (Figure 3).

**Figure 3.**
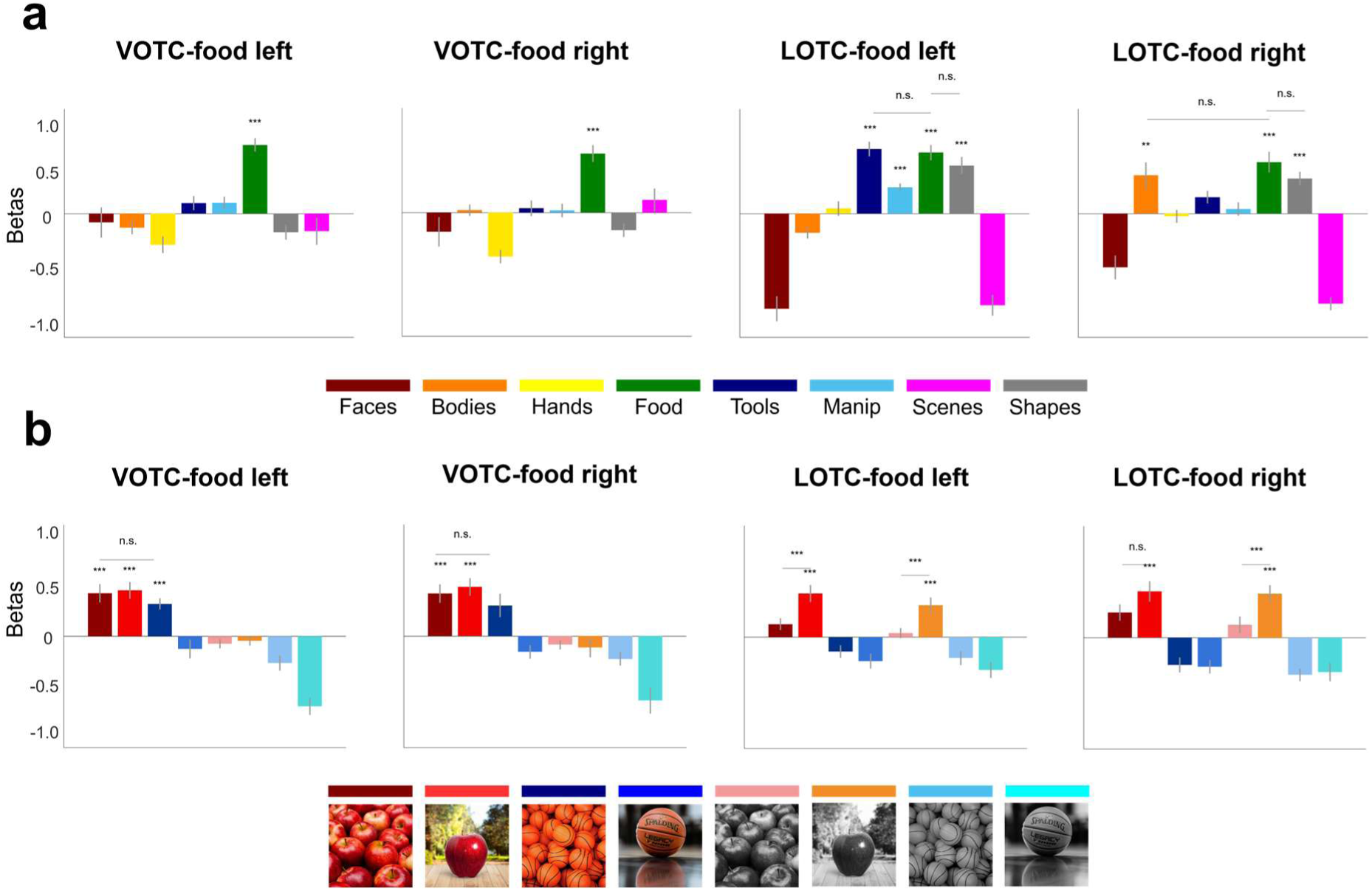
Functional selectivity analysis. Beta values are plotted for each category in food areas identified in each individual participant bilaterally in VOTC and LOTC for **a)** the *category experiment* and **b)** the *surface properties experiment*. Error bars represent ± SEM across subjects (n = 18 participants). Stars (***) represent statistical significance (vs. average of the other categories) Bonferroni corrected at p < .006 (n = 8 comparisons) for the one-sample t-test versus baseline and Bonferroni corrected at p < .007 (n = 7 comparisons) for the pairwise t-test comparisons.

In the *category experiment*, VOTC showed strong selectivity to food with minimal activation to all other categories in both hemispheres (all contrasts p < .001). In contrast, LOTC did not show selectivity for food images, even though its response profile differed based on hemispheric lateralization. In the left hemisphere, responses to food did not differ from tools and meaningless shapes (p > .05), which, together with manipulable objects, all elicited significant higher responses than baseline (all p < .001), as predicted by the left lateralized action gradient. In the right hemisphere, the response to food did not differ from bodies and meaningless shapes (p > .05) and was higher than tools (t_(17)_ = 2.9; p = .01) although this latter difference did not survive correction for multiple comparisons (Bonferroni corrected, n = 7).

In the *surface properties experiment*, VOTC exhibits a complex response pattern driven by the interplay of category preference, colour, and ensemble statistics. A robust activation was observed for coloured food images (p < .001, Bonferroni corrected with n = 8; p = .006) but not for greyscale versions (p > .05). This effect held regardless of whether the stimuli were depicted as single objects or ensembles (both p < .001). Interestingly, inanimate objects also elicited a strong response indistinguishable from food (p > .05) but specifically when presented as coloured ensembles (t_(16)_ = 5.8; p < .001 in the left hemisphere; in the right hemisphere the effect did not survive correction for multiple comparisons; t_(17)_ = 2.6; p = .02; Bonferroni corr. n = 7; p = .007). This high response was not observed for single-coloured objects and all grayscale objects (regardless of format) failed to elicit comparable activation in either hemisphere (all p > .05).

In contrast to VOTC, LOTC food exhibited a response profile driven by object configuration rather than colour. Specifically, while LOTC-food retained a strong preference for food over inanimate objects (all p < .001), this response was not modulated by colour (p > .05). Instead, the region displayed a bias toward single-object processing: responses were higher for single food items compared to ensembles (all p < .006), although this difference did not reach statistical significance in the right hemisphere for coloured conditions (p = .07).

Overall, this pattern suggests a functional dissociation between ventral and lateral food-selective areas in OTC, with the former being sensitive to surface properties of objects, such as colour and ensemble statistics, whereas the latter is tuned to single object configuration but not to colour.

These results were confirmed at the group-level with the vector-of-ROIs analysis, which samples a broad portion of OTC by generating spheres along a vector from the parahippocampal gyrus to the transverse occipital sulcus (see methods). Results are shown in Figure 4.

**Figure 4.**
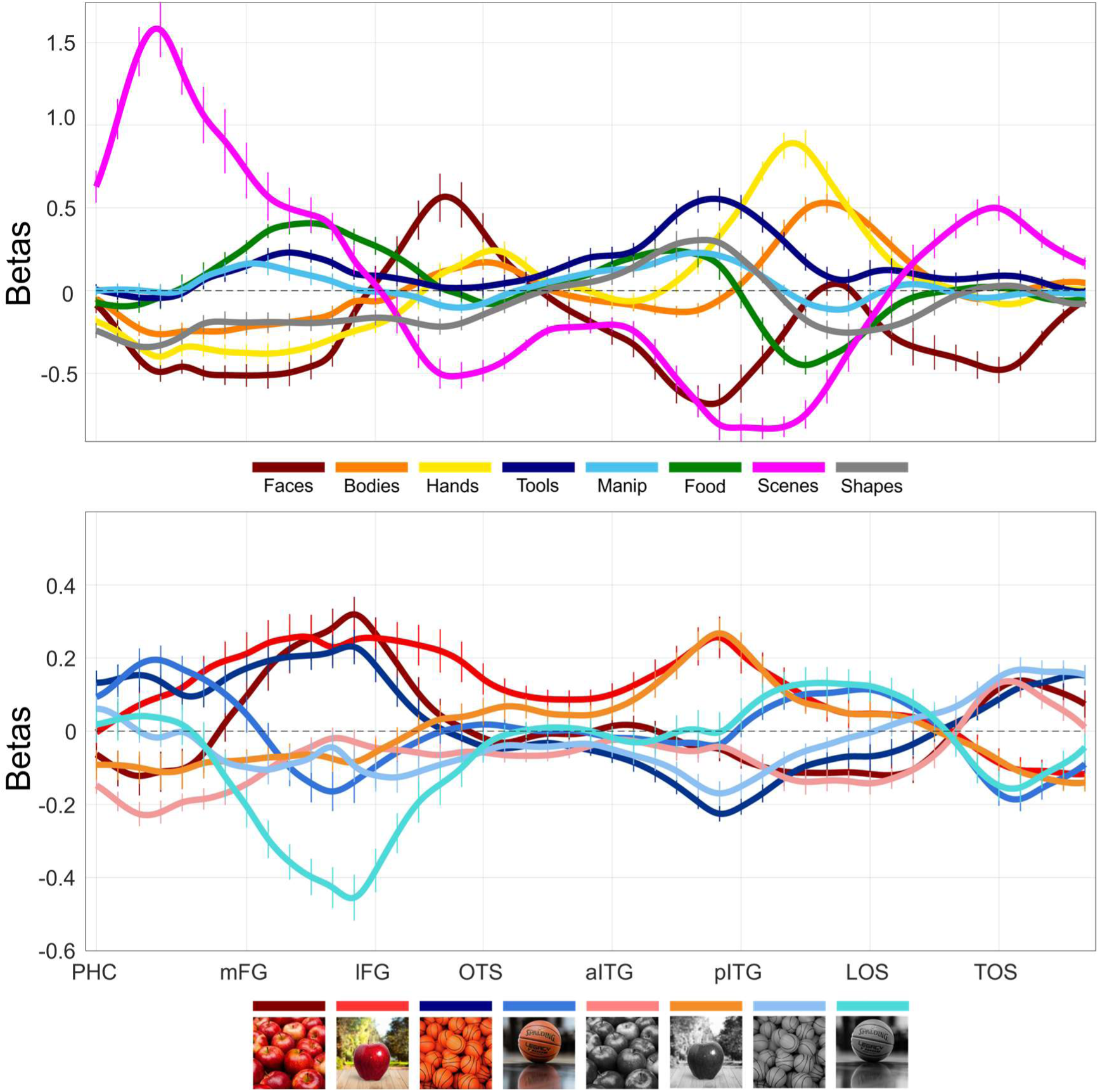
Vector-of-ROIs analysis. A spline connecting distinct anchor points across ventral and lateral OTC is fitted and partially overlapping spheres along the spline are generated. The mean normalized betas for each category are plotted for each sphere. Functional selectivity is visualised for each sphere along the vector (see methods for details). *Top:* vector-of-ROIs for the *category experiment*. Given the relationship found between food, tools, and manipulable objects, we focus on the left hemisphere only. *Bottom*: vector-of-ROIs for the *surface properties experiment*. As no hemispheric differences were found, we averaged across hemispheres. In both plots, the x-axis corresponds to each sphere along the vector, with labels for major anatomical landmarks; the y-axis corresponds to the normalized beta values. Error bars represent ± SEM across participants (n = 18). PHC = Parahippocampal Cortex. mFG = medial Fusiform Gyrus. lFG = lateral Fusiform Gyrus. OTS = Occipitotemporal Sulcus. aITG = anterior Inferior Temporal Gyrus. pITG = posterior Inferior Temporal Gyrus. LOS = Lateral Occipital Sulcus. TOS = Transverse Occipital Sulcus.

Overall, this analysis confirmed the above results. For the *category experiment*, in VOTC, responses to food peaked around the medial fusiform gyrus between scene (parahippocampal cortex) and face (fusiform gyrus) selective cortex, with numerically higher activations compared to inanimate objects. In LOTC, by contrast, the response profile for food follows the same trajectory of other object categories (tools, manipulable objects, and meaningless shapes). Moving posteriorly, response profiles transitioned from food, then tools, to hands and finally to bodies.

For the *surface properties experiment*, VOTC – specifically around the medial fusiform gyrus, which showed strong food responses in the *category experiment* – responded robustly to coloured ensemble stimuli, regardless of whether they depicted food or objects, as well as to images of coloured food presented as single objects. In contrast, LOTC responded preferentially to food, independent of colour, but only when images were presented as single objects, not ensembles.

Taken together, these findings confirm a dissociation between ventral and lateral visual cortex in food specific processing: although both pathways respond to food stimuli, in VOTC, food representations are distinct from other inanimate objects and appear to encode surface properties such as colour and ensemble statistics. On the contrary, in LOTC, food representations are indistinguishable from those of other inanimate objects, likely reflecting action properties of objects. Moreover, LOTC food elicits a response only in a single-object configuration, suggesting a dependence on discrete, graspable shape structure rather than surface cues. Together, these results suggest that the two pathways process distinct information, with the ventral OTC representing primarily surface properties and the lateral OTC sensitive to action-related properties. In the following sections we turn to multivariate analysis to further characterize this dissociation.

### Multivariate analysis reveals the action-related properties underlying food responses

The above results identified food responses in both ventral and lateral OTC but their functional profile differs. Here, we test the role of action-related properties such as manipulability and action effector in driving their representational content. According to the action-based organization reported in our previous study, we predict that food representations should be better characterised by manipulability properties as opposed to action effector properties and therefore showing higher representational similarities with manipulable objects as opposed to tools. To this aim, we computed representational similarity matrices for VOTC and LOTC identified at the group level using the Glasser atlas. For each ROI, we calculated pairwise correlations among the response patterns elicited by the eight stimulus categories, yielding 8 × 8 RSMs. The resulting RSMs for the two ROIs are shown in Figure 5a. Two indices were derived from these matrices (see methods for details). The food (manipulability) index quantifies representational similarities for manipulability properties by measuring the distance between food and other object categories. The hand (action effector) index quantifies the representational similarities for action effector properties by measuring the distance between hands and other object categories. The representational distance for each index was tested with a 2×3 (Region x Category) repeated-measures ANOVA with Region (LOTC, VOTC) and Category (food-manipulable; food-tool, food-hand) as within subject factors. Results confirmed our hypothesis.

**Figure 5.**
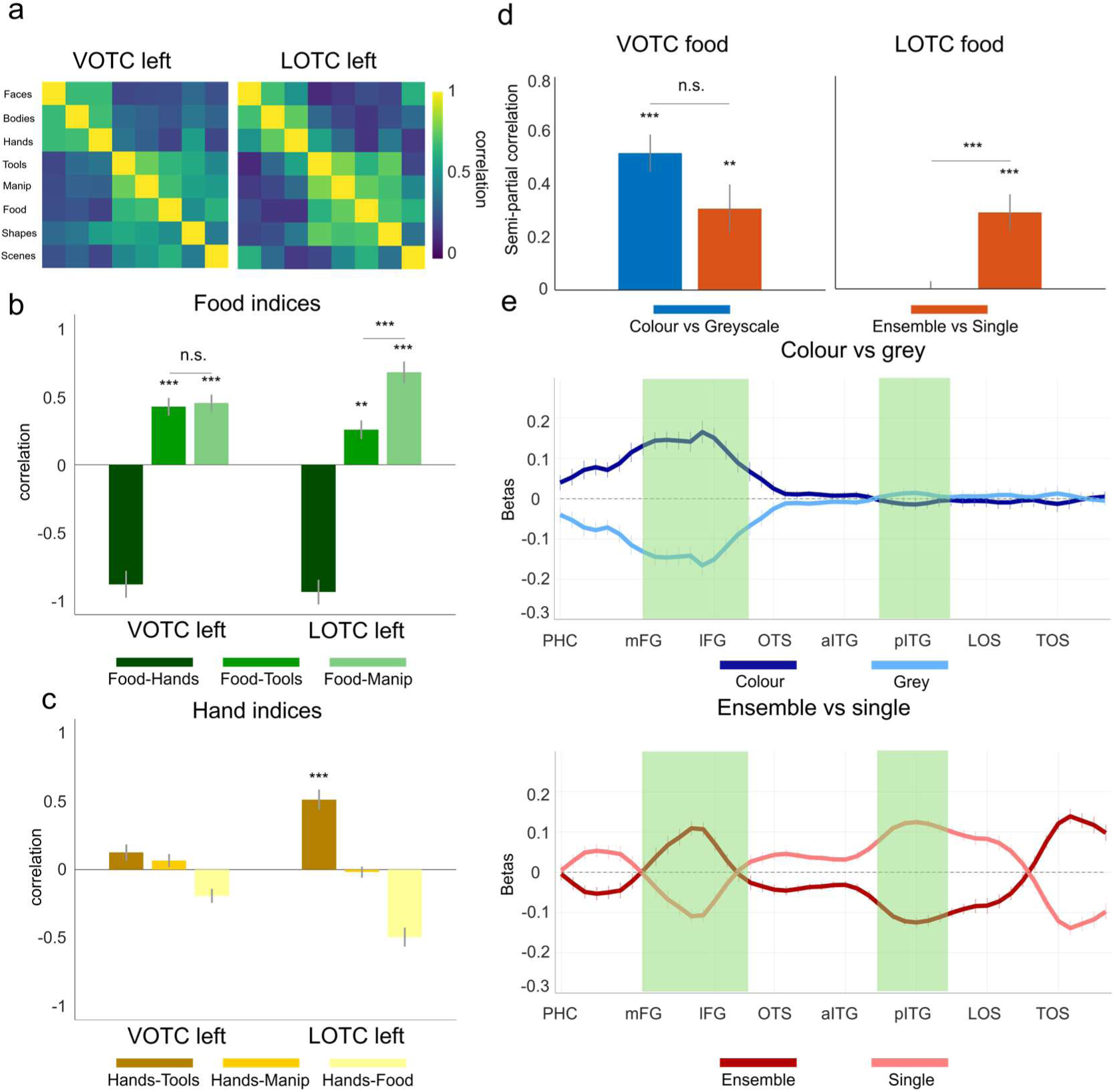
Multivariate analyses reveal dissociation between ventral and lateral OTC. **a)** Correlation matrices for VOTC and LOTC (left hemisphere) defined with the Glasser atlas. The patterns of responses for the eight conditions were correlated with each other to obtain an 8×8 similarity matrix, one for each region. **b)** Food index analysis. The indices indicate the relationship in multivariate space between food and other categories (tools, manipulable objects, and hands). **c)** Hand index analysis. The indices indicate the relationship in multivariate space between hands and other categories (tools, manipulable objects, and food). For both indices, stars (***) represent statistical significance Bonferroni corrected at p < .004 (n = 12 comparisons). **d)** RSA results. Semi-partial correlation was run between the patterns of responses of the ventral and lateral food ROIs (averaged across hemispheres) and two models representing the two surface properties (colour-vs-greyscale and ensemble-vs-single). Stars represent statistical significance Bonferroni corrected with n = 3 comparisons (*** p < .001, and ** p < .01). **e)** Vector-of-ROIs for the two orthogonal dimensions, averaged across hemispheres. For each sphere along the vector, we contrasted the activation for one condition vs. the activation for the other (e.g., ensembles vs single objects). Error bars represent ± SEM across subjects (n = 18 participants). Green boxes underlie the spheres that show a preference for food relative to inanimate objects.

For the food index, the ANOVA revealed a main effect of Category (*F*_(2,34)_ = 92.76, *p* < .001) indicating that food representations show differential similarities with the three action-related categories. Even though the main effect of Region (*F*_(1,17)_ = 3.24, *p* = .089) showed a marginal effect that did not reach significance, the Region × Category interaction was significant (*F*_(2,34)_ = 4.08, *p* = .026). These results suggest that the two regions nevertheless differ in their representational structure. To examine whether the dimensions underlying food are related to manipulability, we conducted post hoc t-tests comparisons. If representational content in these regions reflects manipulability, food should show greater similarity to manipulable objects, which are graspable but not effectors, than to tools, which are both manipulable and associated with effector use. Results confirmed these predictions: LOTC food representations showed significantly higher similarity to graspable objects relative to tools (*t*_(17)_ = 3.60, p = 0.002). On the contrary, in VOTC, food representations were equally similar to both manipulable objects and tools (*t*_(17)_ = 0.33, *p* = 0.74).

These results were further confirmed by the hand index analysis. The 2×3 repeated-measures ANOVA revealed no significant main effect of Region (*F*_(1,17)_ = 1.62, *p* = .221). The main effect of Category was again highly significant (*F*_(2,34)_ = 67.39, *p* < .001). The Region × Category interaction was also significant (*F*_(2,34)_ = 8.93, *p* = .001), demonstrating that the pattern of hand-based distances differed across VOTC and LOTC. Post-hoc paired t-tests revealed a graded pattern: the hand-tool correlation was higher than hand-manipulable correlation (t_(17)_ = 5.46, *p* < .001), which in turn was higher than the hand-food correlation (t_(17)_ = 5.51, *p* < .001). This pattern indicates a representational action gradient. This effect was much weaker in VOTC: although the hand-manipulable correlation was significantly higher than the hand-food correlation (t_(17)_ = 3.24, *p* = 0.005), there was no difference between the hand-tool and the hand-manipulable correlation (t_(17)_ = 0.63, p = 0.53).

These results replicate and extend our previous findings (Cortinovis et al., 2025a), reinforcing the evidence for an action-related organizational gradient: as food is characterised by manipulability properties but not action effector properties, its location in the representational space is closer to that of manipulable objects as opposed to tools which instead more closely relate to hands as they both share action effector properties. This action based representational gradient is observed in lateral but not ventral OTC.

### The role of surface properties in OTC food representations

The functional selectivity analysis revealed that surface properties modulate both VOTC and LOTC in a complex way. To clarify how the two surface properties (colour, ensemble statistics) contribute to the representational structure observed in food-selective cortex, we conducted a Representational Similarity Analysis (RSA). We constructed 8×8 dissimilarity matrices for each of the four food ROIs (identified at the native level in each participant) by computing pairwise dissimilarities across the eight conditions of the surface properties experiment. We then averaged across hemisphere and correlated the resulting neural dissimilarity matrices with two models reflecting the two orthogonal surface properties (colour-vs-grayscale and ensemble-vs-single) using semi-partial correlation to estimate the unique contribution of each dimension while controlling for the other (Figure 5d). To quantify whether the contribution of each model differed across regions and hemispheres, we tested the resulting correlations for each model with a 2×2 repeated-measures ANOVA with Region (VOTC vs LOTC) and Models (colour-vs-greyscale and ensemble-vs-single) as within-subject factors.

Results revealed a significant main effect of Region (*F*_(1,17)_ = 12.64, *p* = .002), while the main effect of Model was not significant (*F*_(1,17)_ = 0.29, *p* = .6). Most importantly, however, the Region × Model interaction was significant (*F*_(1,17)_ = 10.25, *p* = .005), demonstrating that the contribution of colour and ensemble statistics differed across ventral and lateral OTC. Post-hoc t-tests further clarified these patterns: in VOTC, both models did not differ from one another (*t*_(17)_ = 1.75, *p* = .1) and differed from baseline (both contrasts: *t*_(17)_ > 3.00, *p* < .006) suggesting that both properties contribute to the representational structure within this region. In contrast, LOTC showed a significant effect only for ensemble-vs-single model which differ from both baseline (*t*_(17)_ = −0.30, *p* = .768) and from the colour-vs-greyscale model, (*t_(17)_* = −3.79, *p* = .002), while this latter model did not differ from baseline (*t*_(17)_ = −0.30, *p* = .768).

To clarify the direction of the information represented, we visualized responses to each dimension independently along the vector-of-ROIs (e.g., contrasting all coloured vs. grayscale conditions across spheres). This analysis informs on the directionality of the effects reported with the RSA: for example, while an ROI might show a significant effect for the ensemble-vs-single model, indicating a representational difference between ensembles and single objects in this area, this analysis cannot tell us the direction of the effect (e.g., if one area is more sensitive to ensembles or to single objects). Results can be visualized in Figure 5e. They reveal that VOTC food is sensitive to both colour and ensemble statistics properties of objects; viceversa, LOTC food responds exclusively to single objects and is not sensitive to colour information. This pattern aligns with prior demonstrations that VOTC is tuned to surface-based properties such as colour and texture/ensemble statistics (Cavina-Pratesi et al., 2010; Cant and Xu, 2012), while LOTC is more sensitive to shape-defined, single-object structure against a clear background (Grill-Spector et al. 2001; Cavina-Pratesi et al., 2010; Xu et al., 2023).

Together, these analyses demonstrate that ventral food-selective areas are primarily driven by surface-based dimensions such as colour and secondarily ensemble statistics, whereas lateral food areas prefer food in a single object configuration, consistent with the functional dissociation between ventral and lateral OTC reported in the previous functional analyses.

#### Food selectivity in TDANNs is organized into two clusters with distinct functional properties

Finally, we evaluated a model of the spatial organization of ventral visual cortex (the TDANN; Margalit et al., 2024) to test whether biologically inspired pressures, specifically employing wiring-length minimization combined with a self-supervised training objective, can account for the topographic organization and functional profile of food representations. To this end, we performed univariate analyses analogous to those conducted in human visual cortex. Specifically, we presented both stimulus sets (from the *category experiment* and the *surface properties experiment*) to five independent initializations of the TDANN and examined the spatial organization, functional selectivity, and patterns of overlap of food responses in the final “VTC-like” layer, meant to capture the organization of high-level ventral visual cortex.

To specifically characterize the organization of food-related responses, we identified contiguous clusters of food-responsive units using a DBSCAN algorithm (see methods). When these units were defined based on responses to the category experiment, their functional selectivity was assessed across all conditions of the surface properties experiment, and viceversa.

Results are shown in Figure 6a–d. Across all five model initializations, two spatially contiguous clusters of food-responsive units were identified. When we define units with the surface properties experiment (Figure 6a) and test the functional selectivity using the category experiment stimulus set (Figure 6c), both clusters revealed high selectivity to food, which was higher compared to all other categories (p < .0001, tested with 10000 permutations), with no sensitivity to other categories sharing similar action-related properties (i.e., tools or manipulable objects). Importantly, these two clusters exhibited distinct functional responses when localized with the *category experiment* (Figure 6b) and analysed using the *surface properties experiment* stimulus set (Figure 6d). Although both clusters responded in virtually the same way to coloured and greyscale stimuli, they differed markedly in their responses to food-related conditions. Specifically, whereas one cluster responded strongly to food stimuli presented in a single-object configuration, but not to food ensembles, the second cluster responded robustly to food stimuli in both single and ensemble configurations (all p < .0001, tested with 10000 permutations). This pattern resembles the dissociation observed between ventral and lateral food-selective clusters in human visual cortex with respect to their differential sensitivity to ensemble statistics.

**Figure 6.**
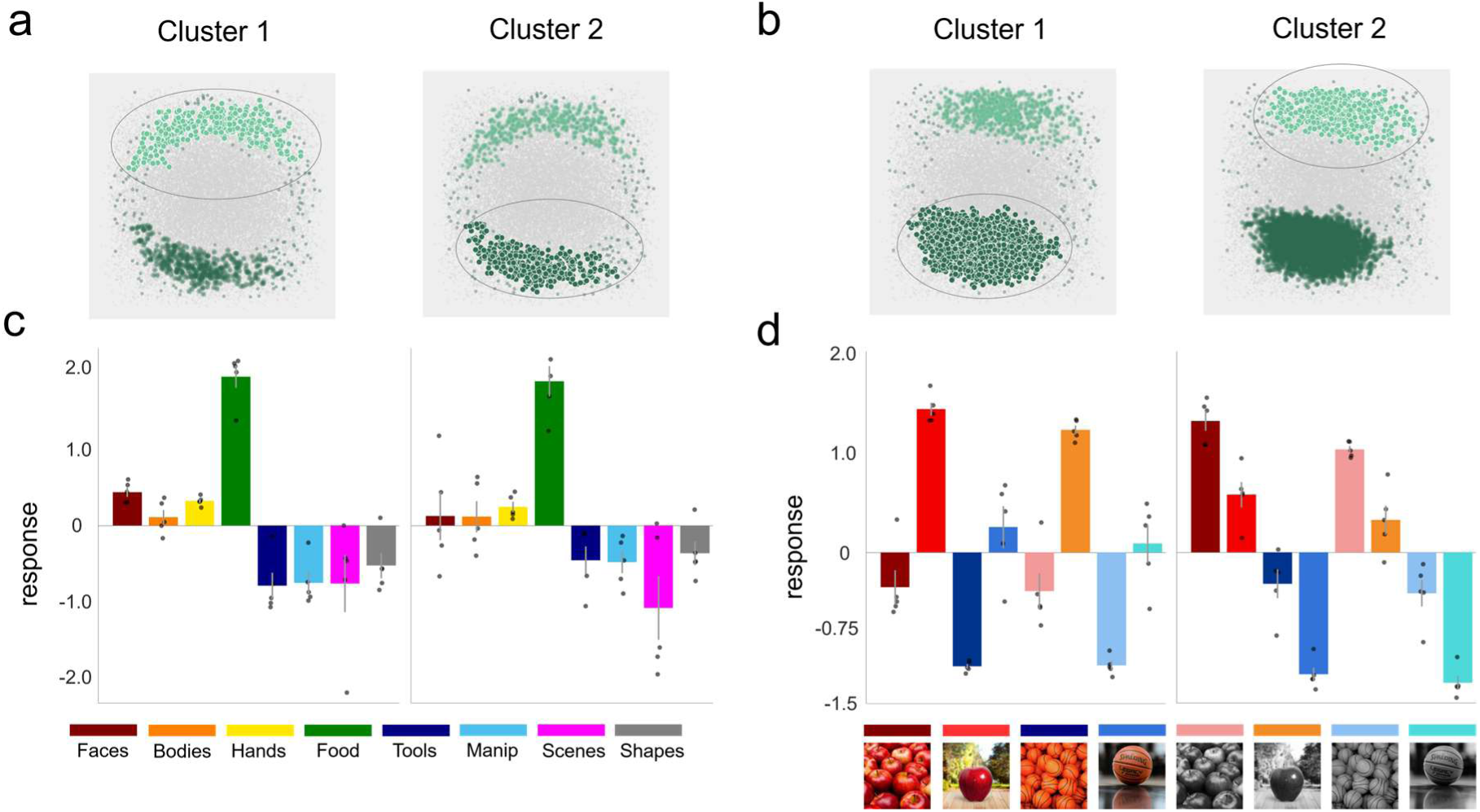
Spatial organization of food selectivity in TDANNs. a) and. **b)** Spatial distribution of food-selective units in the VTC-like layer for one TDANN initialization. Green indicate units selective for food with t > 5.0. Highlighted dots indicate units assigned to a cluster by the DBSCAN algorithm. **c and d)** Selectivity profiles of food-selective units in each identified cluster, based on activations in the VTC-like layer. Responses for each condition are normalized relative to the average response to all other conditions, as in the analyses of human visual cortex. Each data point corresponds to one TDANN model initialization (n = 5). Error bars indicate ± SEM across model initializations.

## Discussion

We investigated the spatial organization, object features, and representational dimensions underlying food representations in ventral and lateral OTC, and found dissociations between the two areas. In lateral OTC, converging analyses indicated that food is not represented as a separate semantic category. Instead, food voxels overlapped with responses to other object categories, the functional profile of this region did not differentiate food from other object categories, and food representations clustered with objects sharing graspable and manipulable properties. As predicted, food representations in this region were more similar to those of manipulable objects than to tools which, beyond being graspable, are also effectors, indicating that multiple action-related dimensions shape object representations in LOTC. In contrast, food voxels in ventral OTC were more strongly associated with visual properties, primarily colour and, to a lesser extent, ensemble statistics. These same properties had little influence on lateral food areas, whose responses were instead tied to single-object configuration. Finally, computational models of OTC spatial organization grouped food responses into two clusters dissociable by sensitivity to ensemble statistics, but could not replicate the colour- or action-related organization observed in visual cortex.

Initial evidence for food-selective areas in human visual cortex came from analyses of the Natural Scenes Dataset (NSD; Allen et al., 2022), which revealed two food-selective clusters: one in ventral OTC near the medial fusiform gyrus and one in lateral OTC around the inferior temporal gyrus. These studies argued against explaining food selectivity through low- or mid-level visual features such as colour, curvature, or texture (Khosla et al., 2022; Jain et al., 2023; Pennock et al., 2023; Henderson et al., 2025). However, the proximity of food-selective areas to known colour-, texture-, and ensemble-responsive regions in ventromedial OTC (Cant and Xu, 2012; Lafer-Sousa et al., 2016), together with reports of overlap between food and tool responses (Ritchie et al., 2024), suggests that food selectivity may be interpreted within the broader spatial organization of multiple feature dimensions mapped in visual cortex (Contier et al., 2024; van Dyck et al., 2026). Using controlled stimulus sets, our study replicates the presence of two food clusters in lateral and ventral OTC and showed that their position and representational content reflect different organizational principles: lateral food selectivity is best understood as part of an action-related gradient, whereas ventral food selectivity relates more closely to regions sensitive to surface-level properties. Below, we consider these two properties in turn.

Previous work reported overlap between food and tool responses in ventral and lateral visual cortex, suggesting that shared action-related properties such as graspability contribute to this organization (Ritchie et al., 2024). We extend this work by showing that action-related properties do explain the emergence of food responses in visual cortex, but primarily in lateral OTC. Only lateral OTC exhibited strong responses to food, tools, and manipulable objects, together with an action-related representational gradient characterized by high similarity between categories sharing action-related properties (hands and tools; food and manipulable objects). These findings are consistent with our proposal that object processing in lateral OTC is organized along an action-related gradient (Cortinovis et al., 2025a), extending from categories associated with end-effector properties (hands and tools) to those associated with general graspability (tools, manipulable objects, and, as shown here, food).

In contrast, food representations in ventral OTC exhibit little-to-no response to tools or manipulable objects, seemingly at odds with previous claims that the medial fusiform gyrus reflects action-related properties of food and tools (Mahon et al., 2007; Ritchie et al., 2024). Instead, our findings suggest that ventral areas preferentially encode surface-level features that relate only indirectly to action and are shared across many inanimate object categories (Mahon and Almeida, 2024; Cortinovis et al., 2025b), including food. This interpretation aligns with proposals of a division of labour between ventral OTC, involved in object processing, and lateral OTC, supporting action processing (Weiner and Grill-Spector, 2013; Lingnau and Downing, 2015; Wurm and Caramazza, 2022). Future work could more precisely identify the graspability- and manipulability-related dimensions organizing food, tools, and other manipulable objects within lateral OTC, extending approaches previously used to characterize action properties of manipulable objects (Almeida et al., 2023; 2025).

Our results demonstrated a strong role for colour in ventral food-selective areas. Colour is especially diagnostic of food: behavioural and neural studies indicate that it supports rapid food detection and recognition more than for many other categories (Conway, 2018; Sato, 2021). Neuroimaging research in humans and macaques consistently identified colour-biased regions in medial OTC, with much weaker colour sensitivity laterally (Brouwer and Heeger, 2009; Lafer-Sousa and Conway, 2013; Lafer-Sousa et al., 2016; Rosenthal et al., 2018; Schalk et al., 2017), and analysis on the NSD similarly revealed correspondence between colour-biased patches and food selectivity (Pennock et al., 2023; 2026). At the same time, studies proposing food selectivity in visual cortex argued that colour cannot fully explain food responses, but relied almost exclusively on the NSD and did not distinguish between ventral and lateral food areas. Using a controlled, hypothesis-driven stimulus set, we show instead that colour is a fundamental property, on par with or exceeding category membership itself, but only in ventral OTC. This novel finding is supported by converging evidence: the overlap between food and colour clusters, similar responses to coloured inanimate objects and non-colour food, and the presence of colour representations in areas identified using only greyscale stimuli.

However, colour is not the only mid-level feature represented in the medial fusiform gyrus/collateral sulcus, which also supports texture and ensemble-statistics processing. In fact, colour-responsive regions in medio-ventral OTC also respond to textures (Cant and Goodale, 2007; 2011; Cavina-Pratesi et al., 2010), and one study identified a region neighbouring and partially overlapping with the parahippocampal place area that is sensitive to ensemble statistics (Cant and Xu, 2012). Importantly, its responses could not be reduced to lower-level texture statistics or colour, but may rely on higher-level texture representations (Cant and Xu, 2015; 2017; Henderson et al., 2023). Here too, a ventral-lateral division emerges: ventral cortex supports texture-like object representations, whereas lateral regions respond more strongly to shape. Our findings support this view by showing that colour and ensemble configuration each explain unique variance in ventral OTC food representations, with colour contributing the most.

Of course, we do not claim that the (left) lateral food cluster is fully explained by action-related properties, nor that colour and ensemble statistics are the only features processed in medio-ventral OTC. Other candidates include shape properties such as curvature and aspect ratio, proposed as organizing principles for both ventral and lateral OTC (Bao et al., 2020; Yue et al., 2020), although these features alone cannot account for food selectivity specifically or category selectivity more generally (Khosla et al., 2022; Yargholi and Op de Beeck, 2023). Similarly, our results cannot distinguish ensemble-specific from high-level texture processing, which likely rely on overlapping mechanisms (Cant and Xu, 2017). Food representations therefore likely reflect a constellation of mid- and high-level properties whose combination is characteristic of food objects.

Analyses of the TDANN, a model designed to capture the spatial organization of visual cortex, revealed both convergence and divergence with neural data. The model exhibited two clusters of food-responsive units that, although not differing in colour sensitivity, diverged in their sensitivity to ensemble statistics: mirroring visual cortex, one cluster responded more strongly to isolated food, whereas the other responded robustly across both configurations, with a bias toward ensembles. Together with our previous work (Cortinovis et al., 2025a), the present results suggest that TDANNs, and ANNs more generally, may better model mid-level visual features than higher-level behaviourally relevant properties (Mahner et al., 2025), including action-related properties. Even colour, often considered a mid-level feature, serves behaviourally relevant goals in humans by signalling potential consumption (Sato, 2021), and is linked to higher-level processing such as semantic memory and language (Zhao et al., 2024; Liu et al., 2025), potentially explaining the absence of differences between colour and greyscale stimuli in the model. More broadly, the central TDANN assumption – that jointly optimizing a self-supervised contrastive objective and a spatial loss is sufficient to reproduce cortical organization (Finzi et al., 2023; Margalit et al., 2024) – is likely incomplete. Capturing the full structure of these pathways may require more biologically plausible training objectives or datasets, such as those approximating the visual experience of children actively exploring their environment (Long et al., 2024; Lu et al., 2026).

In summary, we demonstrate that distinct features drive food selectivity across OTC: action-related properties such as graspability laterally, and surface properties including colour and ensemble statistics ventrally. These findings support the view that category-selective areas reflect underlying behaviourally relevant feature dimensions rather than purely categorical representations (Peelen and Downing, 2017; Bracci and Op de Beeck 2023; Ritchie et al., 2025). Within this framework, food-selective areas are not “food detectors” per se, but regions tuned to visual and semantic properties reliably associated with food and relevant for interacting with it. Which properties are represented depends on each area’s position within a processing pathway and the computational goals that pathway supports.

## Data and code availability

Stimulus sets, MATLAB code used for all analyses, and fMRI data are available on the Open Science Framework at the following link: https://osf.io/27awq

## Acknowledgments

D.C. was supported by a doctoral scholarship provided by the Center for Mind\Brain Sciences at the University of Trento. This work was funded by the Italian Ministry of University and Research (MUR) under the Fondo Italiano per la Scienza – Bando FIS 2, D.D. n. 1236 of 01/08/2023, project code FIS-2023-02203, CUP E53C25001830001, awarded to S.B. Computational resources were provided by the high-performance computing clusters at the University of Trento.

## Notes

https://osf.io/27awq/overview

### Competing Interest Statement

The authors have declared no competing interest.

## References

1. Allen, E. J., St-Yves, G., Wu, Y., Breedlove, J. L., Prince, J. S., Dowdle, L. T., … & Kay, K. (2022). A massive 7T fMRI dataset to bridge cognitive neuroscience and artificial intelligence. Nature neuroscience, 25(1), 116–126.

2. Almeida, J., Fracasso, A., Kristensen, S., Valério, D., Bergström, F., Chakravarthi, R., … & Walbrin, J. (2023). Neural and behavioral signatures of the multidimensionality of manipulable object processing. Communications Biology, 6(1), 940.

3. Almeida, J., Kristensen, S., Tal, Z., & Fracasso, A. (2025). Contentopic mapping in ventral and dorsal association cortex: the topographical organization of manipulable object information. NeuroImage, 121514.

4. Alvarez, G. A. (2011). Representing multiple objects as an ensemble enhances visual cognition. Trends in cognitive sciences, 15(3), 122–131.

5. Arcaro, M., & Livingstone, M. (2024). A whole-brain topographic ontology. Annual Review of Neuroscience, 47.

6. Avery, J. A., Liu, A. G., Ingeholm, J. E., Gotts, S. J., & Martin, A. (2021). Viewing images of foods evokes taste quality-specific activity in gustatory insular cortex. Proceedings of the National Academy of Sciences, 118(2), e2010932118.

7. Bao, P., She, L., McGill, M., & Tsao, D. Y. (2020). A map of object space in primate inferotemporal cortex. Nature, 583(7814), 103–108.

8. Bracci, S., Cavina-Pratesi, C., Ietswaart, M., Caramazza, A., & Peelen, M. V. (2012). Closely overlapping responses to tools and hands in left lateral occipitotemporal cortex. Journal of neurophysiology, 107(5), 1443–1456.

9. Bracci, S., & Op de Beeck, H. P. (2023). Understanding human object vision: a picture is worth a thousand representations. Annual review of psychology, 74(1), 113–135.

10. Bracci, S., & Peelen, M. V. (2013). Body and object effectors: the organization of object representations in high-level visual cortex reflects body–object interactions. Journal of Neuroscience, 33(46), 18247–18258.

11. Brainard, D. H., & Vision, S. (1997). The psychophysics toolbox. Spatial vision, 10(4), 433–436.

12. Brouwer, G. J., & Heeger, D. J. (2009). Decoding and reconstructing colour from responses in human visual cortex. Journal of Neuroscience, 29(44), 13992–14003.

13. Cant, J. S., & Goodale, M. A. (2007). Attention to form or surface properties modulates different regions of human occipitotemporal cortex. Cerebral cortex, 17(3), 713–731.

14. Cant, J. S., & Goodale, M. A. (2011). Scratching beneath the surface: new insights into the functional properties of the lateral occipital area and parahippocampal place area. Journal of Neuroscience, 31(22), 8248–8258.

15. Cant, J. S., & Xu, Y. (2012). Object ensemble processing in human anterior-medial ventral visual cortex. Journal of Neuroscience, 32(22), 7685–7700.

16. Cant, J. S., & Xu, Y. (2015). The impact of density and ratio on object-ensemble representation in human anterior-medial ventral visual cortex. Cerebral Cortex, 25(11), 4226–4239.

17. Cant, J. S., & Xu, Y. (2017). The contribution of object shape and surface properties to object ensemble representation in anterior-medial ventral visual cortex. Journal of cognitive neuroscience, 29(2), 398–412.

18. Cavina-Pratesi, C., Kentridge, R. W., Heywood, C. A., & Milner, A. D. (2010). Separate channels for processing form, texture, and colour: evidence from fMRI adaptation and visual object agnosia. Cerebral cortex, 20(10), 2319–2332.

19. Chen, T., Kornblith, S., Norouzi, M., & Hinton, G. (2020, November). A simple framework for contrastive learning of visual representations. In International conference on machine learning (pp. 1597–1607). PmLR.

20. Contier, O., Baker, C. I., & Hebart, M. N. (2024). Distributed representations of behaviour-derived object dimensions in the human visual system. Nature Human Behaviour, 8(11), 2179–2193.

21. Conway, B. R. (2018). The organization and operation of inferior temporal cortex. Annual review of vision science, 4(1), 381–402.

22. Conway, B. R., Moeller, S., & Tsao, D. Y. (2007). Specialized color modules in macaque extrastriate cortex. Neuron, 56(3), 560–573.

23. Cortinovis, D., Peelen, M. V., & Bracci, S. (2025b). Tool representations in human visual cortex. Journal of Cognitive Neuroscience, 37(3), 515–531.

24. Cortinovis, D., Truong, N., Op de Beeck, H., & Bracci, S. (2025). Investigating action topography in visual cortex and deep artificial neural networks. Nature Communications.

25. Deb, M., Deb, M., & Murty, A. (2025, May). TopoNets: High performing vision and language models with brain-like topography. In International Conference on Learning Representations (Vol. 2025, pp. 39099–39119).

26. Deng, J., Dong, W., Socher, R., Li, L. J., Li, K., & Fei-Fei, L. (2009, June). Imagenet: A large-scale hierarchical image database. In 2009 IEEE conference on computer vision and pattern recognition (pp. 248-255). Ieee.

27. Ester, M., Kriegel, H. P., Sander, J., & Xu, X. (1996, August). A density-based algorithm for discovering clusters in large spatial databases with noise. In kdd (Vol. 96, No. 34, pp. 226–231.

28. Finzi, D., Margalit, E., Kay, K., Yamins, D. L., & Grill-Spector, K. (2023). A single computational objective drives specialization of streams in visual cortex. *bioRxiv*, 2023-12.

29. Foroni, F., Pergola, G., & Rumiati, R. I. (2016). Food color is in the eye of the beholder: The role of human trichromatic vision in food evaluation. Scientific reports, 6(1), 37034.

30. Glasser, M. F., Coalson, T. S., Robinson, E. C., Hacker, C. D., Harwell, J., Yacoub, E., … & Van Essen, D. C. (2016). A multi-modal parcellation of human cerebral cortex. Nature, 536(7615), 171–178.

31. Grill-Spector, K., Kourtzi, Z., & Kanwisher, N. (2001). The lateral occipital complex and its role in object recognition. Vision research, 41(10-11), 1409–1422.

32. Grill-Spector, K., & Weiner, K. S. (2014). The functional architecture of the ventral temporal cortex and its role in categorization. Nature Reviews Neuroscience, 15(8), 536–548.

33. Hasson, U., Harel, M., Levy, I., & Malach, R. (2003). Large-scale mirror-symmetry organization of human occipito-temporal object areas. Neuron, 37(6), 1027–1041.

34. Henderson, M. M., Tarr, M. J., & Wehbe, L. (2023). A texture statistics encoding model reveals hierarchical feature selectivity across human visual cortex. Journal of Neuroscience, 43(22), 4144–4161.

35. Henderson, M. M., Tarr, M. J., & Wehbe, L. (2025). Origins of food selectivity in human visual cortex. Trends in Neurosciences, 48(2), 113–123.

36. Jain, N., Wang, A., Henderson, M. M., Lin, R., Prince, J. S., Tarr, M. J., & Wehbe, L. (2023). Selectivity for food in human ventral visual cortex. Communications Biology, 6(1), 175.

37. Julian, J. B., Fedorenko, E., Webster, J., & Kanwisher, N. (2012). An algorithmic method for functionally defining regions of interest in the ventral visual pathway. Neuroimage, 60(4), 2357–2364.

38. Kanwisher, N. (2010). Functional specificity in the human brain: a window into the functional architecture of the mind. Proceedings of the national academy of sciences, 107(25), 11163–11170.

39. Khosla, M., Murty, N. A. R., & Kanwisher, N. (2022). A highly selective response to food in human visual cortex revealed by hypothesis-free voxel decomposition. Current Biology, 32(19), 4159–4171.

40. Konkle, T., & Caramazza, A. (2013). Tripartite organization of the ventral stream by animacy and object size. Journal of Neuroscience, 33(25), 10235–10242.

41. Konkle, T., & Oliva, A. (2012). A real-world size organization of object responses in occipitotemporal cortex. Neuron, 74(6), 1114–1124.

42. Kriegeskorte, N., Mur, M., & Bandettini, P. A. (2008b). Representational similarity analysis-connecting the branches of systems neuroscience. Frontiers in systems neuroscience, 2, 249.

43. Kriegeskorte, N., Mur, M., Ruff, D. A., Kiani, R., Bodurka, J., Esteky, H., … & Bandettini, -P. A. (2008a). Matching categorical object representations in inferior temporal cortex of man and monkey. Neuron, 60(6), 1126–1141.

44. Lafer-Sousa, R., & Conway, B. R. (2013). Parallel, multi-stage processing of colours, faces and shapes in macaque inferior temporal cortex. Nature neuroscience, 16(12), 1870–1878.

45. Lafer-Sousa, R., Conway, B. R., & Kanwisher, N. G. (2016). Colour-biased regions of the ventral visual pathway lie between face-and place-selective regions in humans, as in macaques. Journal of Neuroscience, 36(5), 1682–1697.

46. Lingnau, A., & Downing, P. E. (2015). The lateral occipitotemporal cortex in action. Trends in cognitive sciences, 19(5), 268–277.

47. Liu, B., Wang, X., Wang, X., Li, Y., Han, Y., Lu, J., … & Bi, Y. (2025). Object knowledge representation in the human visual cortex requires a connection with the language system. PLoS biology, 23(5), e3003161.

48. Long, B., Sparks, R. Z., Xiang, V., Stojanov, S., Yin, Z., Keene, G. E., … & Frank, M. C. (2024). The BabyView dataset: High-resolution egocentric videos of infants’ and young children’s everyday experiences. *arXiv preprint arXiv:2406.10447*.

49. Lu, Z., Doerig, A., Bosch, V., Krahmer, B., Kaiser, D., Cichy, R. M., & Kietzmann, T. C. (2025). End-to-end topographic networks as models of cortical map formation and human visual behaviour. Nature Human Behaviour, 1–17.

50. Lu, Z., Thorat, S., Cichy, R. M., & Kietzmann, T. C. (2026). Adopting a human developmental visual diet yields robust and shape-based AI vision. Nature Machine Intelligence, 1–14.

51. Mahner, F. P., Muttenthaler, L., Güçlü, U., & Hebart, M. N. (2025). Dimensions underlying the representational alignment of deep neural networks with humans. Nature Machine Intelligence, 7(6), 848–859.

52. Mahon, B. Z., & Almeida, J. (2024). Reciprocal interactions among parietal and occipito-temporal representations support everyday object-directed actions. Neuropsychologia, 198, 108841.

53. Mahon, B. Z., Milleville, S. C., Negri, G. A., Rumiati, R. I., Caramazza, A., & Martin, A. (2007). Action-related properties shape object representations in the ventral stream. Neuron, 55(3), 507–520.

54. Margalit, E., Lee, H., Finzi, D., DiCarlo, J. J., Grill-Spector, K., & Yamins, D. L. (2024). A unifying framework for functional organization in early and higher ventral visual cortex. Neuron, 112(14), 2435–2451.

55. Matić, K., de Beeck, H. O., & Bracci, S. (2020). It’s not all about looks: The role of object shape in parietal representations of manual tools. Cortex, 133, 358–370.

56. Op de Beeck, H. P., Baker, C. I., DiCarlo, J. J., & Kanwisher, N. G. (2006). Discrimination training alters object representations in human extrastriate cortex. Journal of Neuroscience, 26(50), 13025–13036.

57. Peelen, M. V., & Downing, P. E. (2017). Category selectivity in human visual cortex: Beyond visual object recognition. Neuropsychologia, 105, 177–183.

58. Pennock, I. M., Racey, C., Allen, E. J., Wu, Y., Naselaris, T., Kay, K. N., … & Bosten, J. M. (2023). Colour-biased regions in the ventral visual pathway are food selective. Current Biology, 33(1), 134–146.

59. Pennock, I. M., Racey, C., Kay, K., Franklin, A., & Bosten, J. M. (2026). The colors of images preferred by individual voxels can be used to delineate functionally distinct visually responsive brain areas. Proceedings of the National Academy of Sciences, 123(15), e2535986123.

60. Ritchie, J. B., Andrews, S. T., Vaziri-Pashkam, M., & Baker, C. I. (2024). Graspable foods and tools elicit similar responses in visual cortex. Cerebral Cortex, 34(9), bhae383.

61. Ritchie, J. B., Wardle, S. G., Vaziri-Pashkam, M., Kravitz, D. J., & Baker, C. I. (2025). Rethinking category-selectivity in human visual cortex. Cognitive neuroscience, 1–28.

62. Rosenthal, I., Ratnasingam, S., Haile, T., Eastman, S., Fuller-Deets, J., & Conway, B. R. (2018). Colour statistics of objects, and colour tuning of object cortex in macaque monkey. Journal of vision, 18(11), 1–1.

63. Sato, W. (2021). Colour’s indispensable role in the rapid detection of food. Frontiers in Psychology, 12, 753654.

64. Schalk, G., Kapeller, C., Guger, C., Ogawa, H., Hiroshima, S., Lafer-Sousa, R., … & Kanwisher, N. (2017). Facephenes and rainbows: Causal evidence for functional and anatomical specificity of face and color processing in the human brain. Proceedings of the National Academy of Sciences, 114(46), 12285–12290.

65. Simmons, W. K., Martin, A., & Barsalou, L. W. (2005). Pictures of appetizing foods activate gustatory cortices for taste and reward. Cerebral cortex, 15(10), 1602–1608.

66. Teghipco, A. (2023). Brainsurfer. Zenodo.

67. van der Laan, L. N., De Ridder, D. T., Viergever, M. A., & Smeets, P. A. (2011). The first taste is always with the eyes: a meta-analysis on the neural correlates of processing visual food cues. Neuroimage, 55(1), 296–303.

68. van Dyck, L. E., Hebart, M. N., & Dobs, K. (2026). Multidimensional feature tuning in category-selective areas of human visual cortex. Journal of Neuroscience.

69. Weiner, K. S., & Grill-Spector, K. (2013). Neural representations of faces and limbs neighbor in human high-level visual cortex: evidence for a new organization principle. Psychological research, 77(1), 74–97.

70. Whitney, D., & Yamanashi Leib, A. (2018). Ensemble perception. Annual review of psychology, 69(1), 105–129.

71. Wurm, M. F., & Caramazza, A. (2022). Two ‘what’ pathways for action and object recognition. Trends in cognitive sciences, 26(2), 103–116.

72. Xu, Y., Vignali, L., Sigismondi, F., Crepaldi, D., Bottini, R., & Collignon, O. (2023). Similar object shape representation encoded in the inferolateral occipitotemporal cortex of sighted and early blind people. PLoS Biology, 21(7), e3001930.

73. Yargholi, E., & Op de Beeck, H. (2023). Category trumps shape as an organizational principle of object space in the human occipitotemporal cortex. The Journal of Neuroscience, 43(16), 2960–2972.

74. Yue, X., Robert, S., & Ungerleider, L. G. (2020). Curvature processing in human visual cortical areas. NeuroImage, 222, 117295.

75. Zhao, M., Xin, Y., Deng, H., Zuo, Z., Wang, X., Bi, Y., & Liu, N. (2024). Object color knowledge representation occurs in the macaque brain despite the absence of a developed language system. PLoS Biology, 22(10), e3002863.

